# Using Aptamers for Protein Scale-up

**DOI:** 10.1101/2025.03.04.641353

**Authors:** Andrea Callegaro, Xinran Peng, Flemming Morsch, Christian Code

## Abstract

Purifying and concentrating proteins is fundamental to biomedical research for both diagnostics and therapeutics purposes. Several methods can be used for protein purification, ranging from simple chemical methods to more advanced chromatography techniques. We looked at improving research solutions for analysing and concentrating proteins using DNA aptamers for specific soluble proteins. Aptamers can be combined with conventional methods like affinity chromatography by attaching an aptamer to silica or magnetics beads. The protein is bound and subsequently eluted from the bead to yield purified protein.

We have developed computational approaches for developing aptamers against any protein which eliminates the need for making a recombinant protein with a tag, allowing for the purification of more native proteins. In this study a computational approach was used to determine the binding sites between an existing reference aptamer (R_apt) to a well characterised protein, Bovine Serum Albumin (BSA). We found that R_apt binds to BSA specifically at domains I and III. Following this we characterised the binding of R_apt to BSA in vitro using the intercalating dye, SYBR Green I, to show a dissociation constant (K_D_) 0.02 µM. The R_apt was modified by adding 5 adenosines to the 5’ end to make a polyA tail. This aptamer with a modified polyA tail, named Ni_apt, allows binding to Ni-NTA magnetic beads. The Ni_apt had a dissociation constant (K_D_) to R_apt to be 0.12 µM. Lastly, we utilised Ni-NTA magnetic beads coupled with the Ni_apt aptamer to bind and purify BSA from a concentrated solution. We recovered 20.7% of the BSA using our protocol. In future developments, we aim to extend our technology based on this foundation to target proteins with therapeutic or diagnostic potential, such as extracting and concentrating immunoglobulins, antibodies and high value proteins.

## 1. Introduction

Aptamers are single stranded non-coding nucleic acid (DNA and RNA) molecules consisting of 10-140 nucleic acid bases that fold into 3D structures (Ellington, 1990). Aptamers are also known as chemical antibodies and are optimised for selection against a variety of targets including proteins and small molecules (McKeague, 2012) as well as a number of cells (Hamula et al. 2008) and viruses (Sánchez-Báscones et al. 2021). Developing aptamers against specific protein biomarkers relevant to specific diseases can provide a novel approach to diagnosing diseases especially when there is not a suitable antibody available (Dunn et al., 2017). Specific aptamers have been used to identify serum biomarkers for diseases like liver fibrosis (Luo, et al 2021) and cancer (Tang et al. 2007; Rom et al. 2011).

Although aptamers can act on similar targets as antibodies, they significantly differ from them. Aptamers do not cause strong innate immunogenicity, are chemically synthesised, more stable, and do not have batch to batch variability. A table of the comparisons can be found in Ali review (2019). Aptamers have until recently been selected through SELEX methods (Tuerk & Gold 1990) to find an aptamer against a specific target. Several computational methods have allowed aptamers to be designed in silico and a review can be found by Sun et al (2022). Recently, Large Language Models (LLM) have been used to predict better aptamer sequences (Morsch et al. 2023) based on their physicochemical properties and create 2D (Nussinov et al 1980; Zuker et al 1981; Zuker 2003; SantaLucia 2004) and 3D rendering (Yi et al 2022). The 3D rendered aptamer can be used to dock the aptamer to its cognate protein to outline potential specific regions where the aptamer binds (Seperi Zarandi et al. 2020; Jimenez-Garcia et al. 2023). These in silico tools both allow a faster way to develop and visualise aptamers binding to their targets. Finally, the dissociation constant (Kd) can be determined using SYBR Green I, a forced Intercalation probe (Fig. 1) which has been used extensively for aptamer research (McKeague, 2012; 2015).

**Figure 1.**
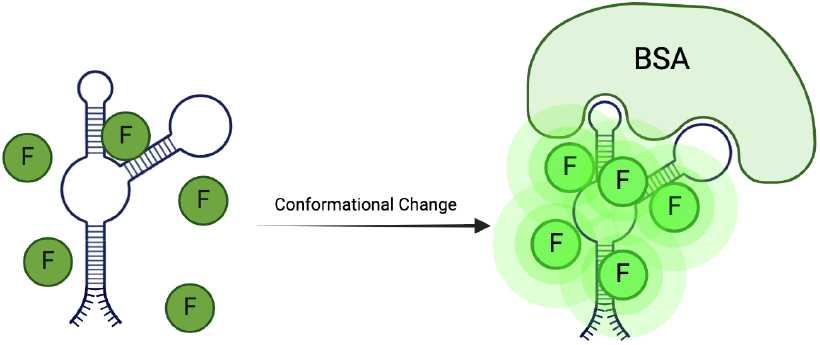
Cartoon of SYBR Green I binding to aptamer and release from aptamer in absence (left panel) and presence (right panel) of target. Note this picture could be shown the opposite way where SYBR Green I is intercalated into the aptamer with more stacked character.

Besides its diagnostic and therapeutic uses, aptamers can be employed to upconcentrate proteins, which overcomes the drawbacks of traditional techniques for protein purification such as ion-exchange, size-exclusion and affinity chromatography (Zhao et al. 2012). These techniques require a lot of work, specific tags and present limitations: non-homogeneous dispersion, high cost and scalability issues for industrial application (Hamdan et al. 2002; Aguilar, 2004; Ramos-de-la-Pena et al 2019; Liu 2020). In this study we focused on upconcentrating and purifying a well studied and inexpensive protein, BSA, using aptamer-based affinity chromatography. BSA was used for both pragmatic and cost efficiency reasons, but the methods presented can be applied to more expensive and complex proteins. This method could be used as a novel way to concentrate concentrated glycated albumin to determine diabetes (Freitas, 2017) or else, study the binding interactions among serum protein and cancer biomarkers (Cortez, Célia Martins, et al 2012).

In this research the BSA aptamer (R_apt) was originally selected using SELEX methods (Wang et al 2019). We added upon these initial studies to develop a 3D rendering and dock R_apt to the BSA protein and found binding sites. We found that the R_apt had conserved binding regions within the major loop of the aptamer. Since the 5’ end was not involved in the binding we added a polyA tail tag to couple it to Ni-NTA magnetic beads (Nastasijevic et al. 2008). The process is based on BSA attachment to magnetic bead-bound aptamers and the subsequent measurement of the final concentration of unbounded BSA (Fig. 2). This new aptamer-based chromatography optimises the traditional affinity chromatography with high efficiency, specificity and low cost to upconcentrate specifically BSA and possibly other proteins. We found that the addition of the polyA tail did not affect the binding affinity of the aptamer to the BSA, even if the computational predictions showed a change in aptamer folding and binding region of BSA protein. The R_apt had a Kd of 0.10 µM and 0.16 µM for the Ni_apt. We utilised the Ni_apt to up-concentrate the BSA quickly and easily. We found that the aptamer bound magnetic beads functionally and was able to bind 20.7 % of the protein in the sample. Overall, this study shows an efficient method to modify an aptamer to be used for protein purification.

**Figure 2.**
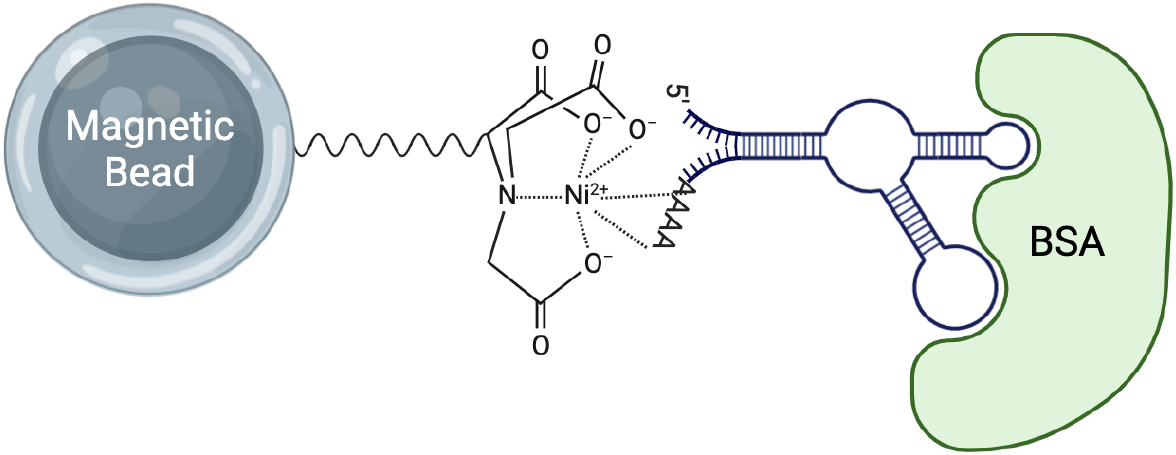
Representation of the Ni_apta-Based Affinity Chromatography system for up-concentrating BSA using Ni-NTA magnetic beads. Ni-NTA magnetic beads treated with Ni_apt allow the functional binding of BSA to the matrix mimicking the His tag and leaving the elute out, upconcentrating BSA protein in the solution.

## 2. Materials and Methods

### 2.1 Chemicals and Reagents

The sequences of the aptamers used are:

R_apt-5’ - AGCAGCACAGAGGTCAGATGGTATCGAGCGCAGGGCCGCCTTTGTT-3’; Ni_apt-5’-AAAAAAAAA AGCAGCACAGAGGTCAGATGGTATCGAGCGCAGGGCCGCCTTTGTT-3’ (Merck, Haverhille, UK). Bovine Serum Albumin: probumin, bovine serum albumin, diagnostic grade (Millipore, Il, USA). His-Mag beads Ni-NTA beads (Cytivia, SE). Binding Buffer was used for all experiments (100 mM NaCl, 20 mM Tris–HCl, 2 mM MgCl_2_, 5 mM KCl, 1 mM CaCl_2_, pH 7.6) (Merck, MO, USA). All salts Calcium Chloride, Magnesium Chloride, Sodium Chloride and Tris (Merck, MO, USA). SYBR Green I (Thiazole Green) in 10,000x in DMSO (Biotium, CA, USA) (10,000× 10 mg/ml, 19.6 mM with a molar absorption coefficient of 73000 M-1 cm-1 in buffer) (Zipper et al. 2004) was diluted to 1x in binding buffer by adding 10 µl of SYBR Green I to 100 mL of binding buffer. Imidazole powder (Merck, MO, USA) was diluted in the binding buffer to obtain a solution of 10 mM used to regenerate the magnetic beads.

### 2.2 Aptamer 2D & 3D Models and Docking

The 2D and 3D models of R_apt and Ni_apt were prepared using Unafold and 3dRNA/DNA setting the ion concentrations based on the buffer utilised in the study of Zhang et al (2022 a&b). Docking was performed with HDOCK (Huang et al. 2014; Yan et al. 2017) and AutoDock/Vina (Seeliger et al 2010) using the BSA (pdb:4F5S) as the protein input (Bujacz, 2012). The 3D model and docking was visualised in Pymol (Schrodinger, NY, USA) (Schrödinger et al. 2020), while the study of binding interactions was performed with PLIP (Adasme et al. 2021).

### 2.3 Aptamer Preparation

Both aptamers were diluted in the binding buffer to a 100 µM concentration. The solution was heated above the melting temperature 96.2°C for 10 minutes to denature secondary structures, then cooled on ice for 5 minutes. The aptamers were allowed to equilibrate at room temperature for at least 2 hours before use. Before their use the aptamers were further diluted to 10 µM.

### 2.4 Protein Preparation

BSA powder was weighed and dissolved in the Tris binding buffer. The concentration of the resulting BSA (MW = 66,463 g/mol) solution was 12.4 mM. The molar extinction coefficient of BSA at 280 nm is 43,824 M-1 cm-1. The protein concentration was measured with a Thermo Scientific Nanodrop spectrophotometer (Waltham, MA) to ensure correct calculations and avoid weight errors.

### 2.4 Fluorescence Measurements for Aptamers-BSA Binding Interactions

#### 2.4.1 Binding Interactions of R_apt or Ni_apt with BSA

Initially, the concentration of BSA was aliquoted as different working volumes, creating substantial variations during measurements, the following protocol is the optimised version with reduction of operation errors. An aliquot of 1 μL of R_apt or Ni_apt was incubated with 23 μL of SYBR Green I 1x solution (SYBR Green I in Tris binding buffer), in a microplate shaker at room temperature for 5 minutes; fluorescence (F0) was measured with a BMG LabTech Fluorostar Omega spectrophotometer (Germany). Subsequently, 1 μL of BSA protein concentrated solutions were added to the same wells and incubated in a microplate shaker at room temperature for 5 minutes, followed by fluorescence measurement to obtain F1. The final solution volume was 25 μL, with a final aptamer concentration of 0.4 μM. The final protein concentrations were 11.3, 10, 8, 5.65, 4, 2, 0.4, 0.2, 1, 0.04 and 0 μM for the control. The excitation and emission wavelengths were set at 485 nm and 525 nm. Generally, SYBR Green I was utilised to monitor the binding and conformational changes in the aptamers upon binding to BSA. When the aptamers bind to BSA, SYBR Green I is either released or absorbed to the aptamer due to the change in double stranded portions of DNA. For the R_apt and Ni_apt binding to BSA the latter happened as illustrated in Figure 1. As a result, the aptamer-protein interaction can be quantitatively assessed by measuring the fluorescence intensity of SYBR Green I. The aptamer - SYBR Green I ratio was maintained using the method described by Radhika et al. (2021) and Kong et al. (2013), with adjustments made to exceed the nucleotide to dye ratio (Zipper et al. 2004; Radhika et al 2021).

The resultant fluorescence was calculated using:

**(F1-F0) /F0*100 = Relative Delta %**

The Relative Delta % was plotted against the BSA concentration.

### 2.5 Calculation of Dissociation Constant (K_D_) Values and Maximum Binding Affinity Values (B_max_)

Dissociation constant (Kd) and maximum binding capacity (Bmax) were calculated based on methods mentioned in previous studies (Wang *et al*., 2019; Jarmoskaite et al., 2020), with the following the formula:

**Bound = ?max · [Ligand]/(Kd + [Ligand])**

The percentage change in fluorescence is related to the binding of BSA to aptamers, thus was used to calculate the Kd and Bmax in accordance with the fitting model shown in figure 5. (Calculations were based on data fitting curves performed using SciPy, Phyton)

### 2.6 Extraction of BSA using Aptamer Functionalized Magnetic Beads

Beads with Ni-NTA were purchased from Cytiva (Uppsala, Sweden). An aliquot of 100 µl of bead slurry was added to an Eppendorf tube (Eppendorf, Germany). Three washes of 200 µl of imidazole 10 mM solution were used to regenerate the beads and ensure their functionality. Subsequently, five washes were performed with 200 µl of binding buffer to eliminate traces of imidazole. The Ni_apt (1 µl of 10 µM) was added to the beads and was incubated for 30 minutes at room temperature in a tube rotator (speed 30 rpm). The excess aptamer solution was removed to prevent any DNA contamination in the further analysis. A 200 µl aliquot of BSA solution (121 µM) was added to the aptamer complexed beads, after removal of the elute, and incubated for 30 minutes at room temperature in the tube rotator (speed 30 rpm) (Thermo Fischer, MA, USA). The elute was taken and measured with the Nanodrop to measure the unbound BSA. BSA bound to aptamers-beads complex was calculated based on BSA initial concentration (121 µM), final solution volume (200 µl) and eluate final concentration measured with the Nanodrop (Thermo Fischer, MA, USA). The same protocol was repeated for 150 µl and 200 µl beads slurry, maintaining the same final volumes concentrating the beads in the solution and increasing the aptamer volume to 1.5 µl.

#### Calculation of bead Total Binding Capacity = 50mg/mL×0.025mL=1.25mg

Considering our Ni_apt concentration of 0.099 µM, the amount of added DNA (171 µg) functionalized part of the available surface of the beads. We calculated the available beads surface considering the volume of slurry and beads specifications, showing a total surface area for binding of approximately 3.75 billion µm^2^ based on 746,380 beads present in the slurry.

## 3. Results

### 3.1 Selection and Structures of Aptamers

The sequence of the R_apt used in our study (5’-AGCAGCACAGAGGTCAGATGGTATCGAGCGCAGGGCCGCCTTTGTT-3’) was designed based on the findings of Wang et al. developed a label-free and modification-free in situ selection strategy for identifying BSA-specific aptamers. This method utilised an electrophoretic mobility shift assay to screen aptamers, which allows full exposure of single-stranded DNA sequences to interact with BSA that is pre-mixed with the agarose gel (Wang et al., 2019). BSA was a pragmatic and inexpensive protein to develop an aptamer upconcentrating method. We used a polyA tail as it offered a cheaper alternative to adding streptavidin or a His-Tag linker and would reduce nonspecific binding from these tags. A polyA tail on the aptamer allowed it to bind simply to Ni-NTA magnetic beads without adding any additional modification at the 5’ end. The sequence was only extended by 10 nb which was initially thought not to change the structure of the aptamer. The Ni_apt sequence is based on our standard R_apt sequence (5’-AAAAAAAAAAGCAGCACAGAGGTCAGATGGTATCGAGCGCAGGGCCGCCTTTGTT-3’) with the polyA tail added to the 5’ end which mimics a His-Tag (Nastasijevic 2008; Jahan 2023).

The predicted 2D structure (Fig. 1A left panel R_apt; Fig. 1A right panel Ni_apt) and 3D structure (Fig. 1B left panel R_apt; Fig. 1B right panel Ni_apt) of the aptamers used in this study were obtained using Unafold and 3dDNA. The 2D structure forms an internal loop for the BSA aptamer and a bulge for the polyA BSA aptamer. The 3D structure (Fig 3B) shows similar structures with different orientation for both aptamers, inferring to different binding sites on BSA. As BSA is well characterised, several crystal structures of BSA are available through the PBD server, however we chose PDB ID: 4F5S because it was the full protein and had a high resolution of 2.47 Å. The aptamers were docked on the BSA using the online tool HDOCK (Fig 3C) using similar buffer conditions (Yan et al 2017). The prediction was set to select the lowest free energy binding patterns, which reflects optimum binding efficiency and stability. The interactions within the complex were analysed and measured using PyMol and PLIP, and the results were compared with existing literature to validate our findings. Both aptamers bind to similar regions on the BSA protein, particularly domain I and III, and the polyA tail does not change where the aptamer interacts with the protein. The polyA tail therefore can be used for protein purification without interfering with the binding capacity. More detailed binding interactions are discussed in section 3.2 and 4.2.

**Figure 3.**
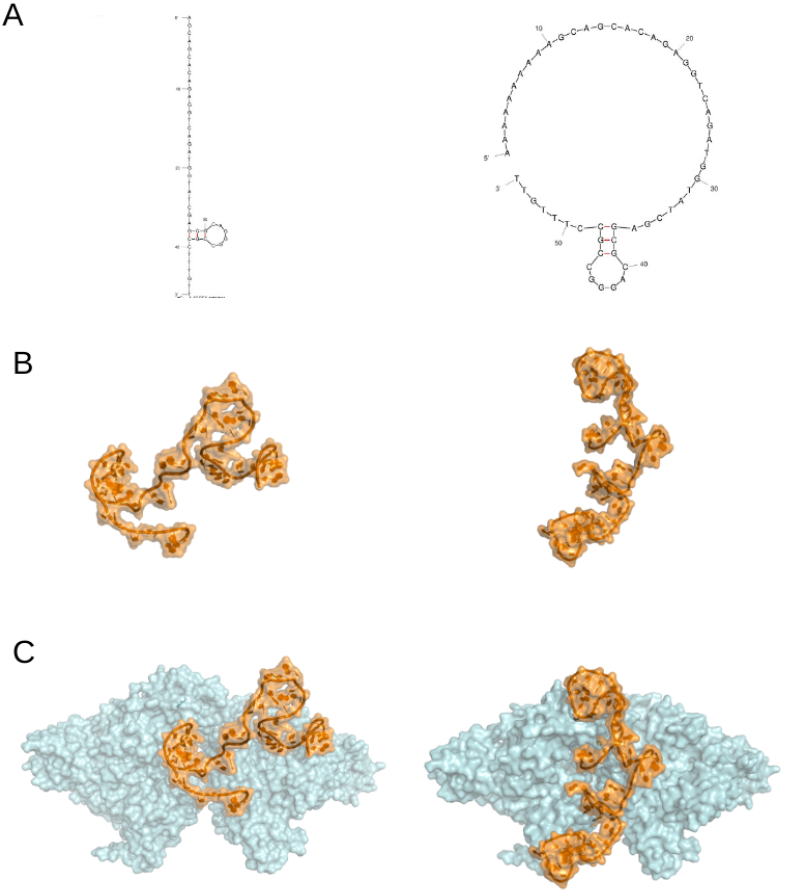
Structure and docking predictions. (A) Predicted Minimum Free Energy (MFE) 2D Structure encoding positional entropy of the R_apt (*left panel*) and the Ni_apt (*right panel*). (B) Predicted 3D structure of the R_apt and Ni_apt shown in orange. (D) Docking prediction between BSA asymmetric unit (cyan) and both aptamers (orange) with minimum free energy. Outlined by the docking predictions the two different aptamers Images in panel B and C were created using PyMol (Schrödinger et al. 2020).

### 3.2 Aptamer-Protein Interactions

We found that the aptamers docked to BSA with a high degree of confidence. The several interaction points between the docked aptamer and the protein were further analysed. PLIP (Adasme et al. 2021) was used to identify and label the interaction points between the aptamer and the protein. As displayed in Fig. 4 and Table 1, we found both aptamers to bind via hydrophobic interactions, hydrogen bonds and salt bridges with both chains of BSA.

**Table 1.**
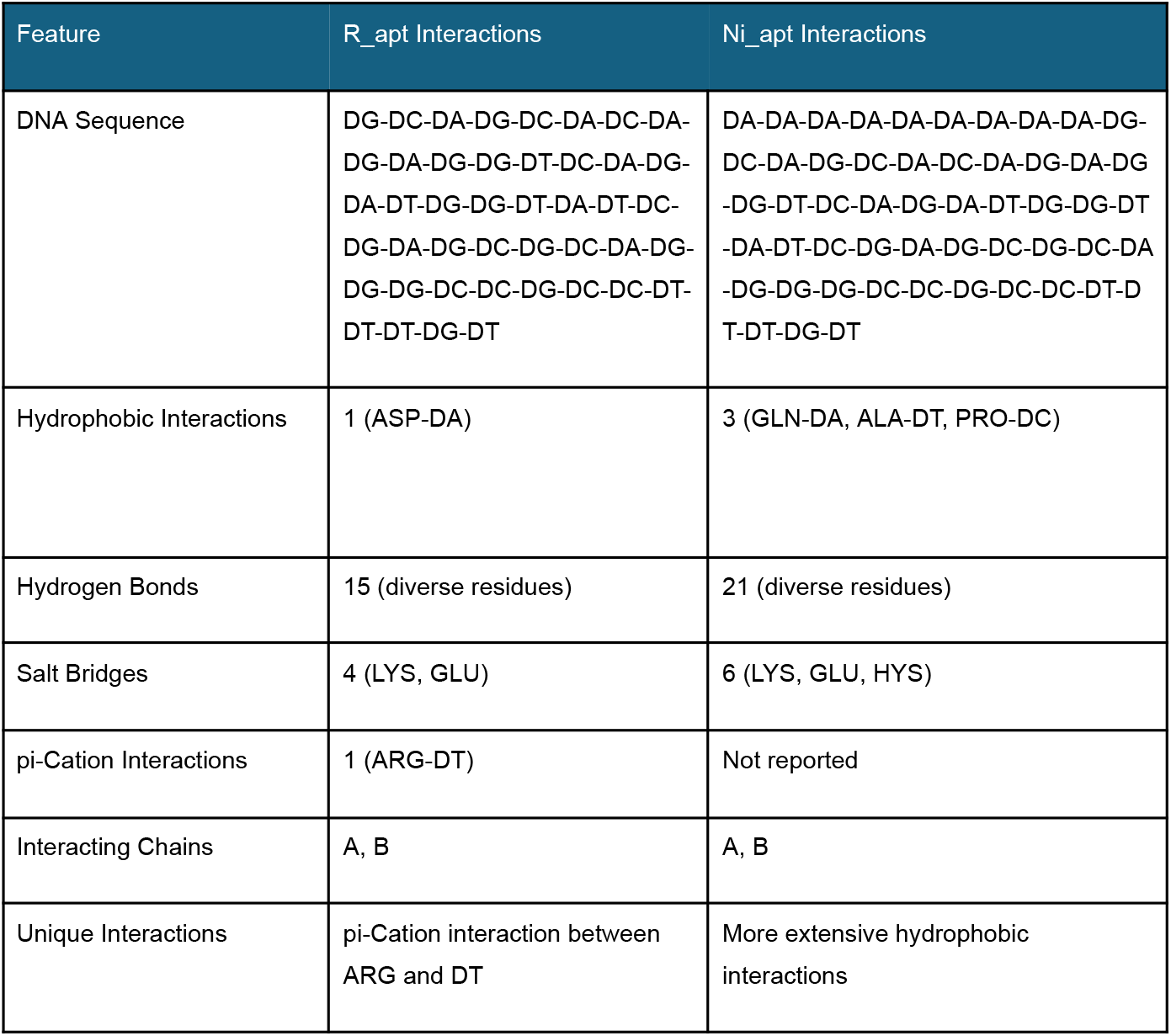
BSA R_apt and BSA Ni_apt Interaction Sites.

**Figure 4.**
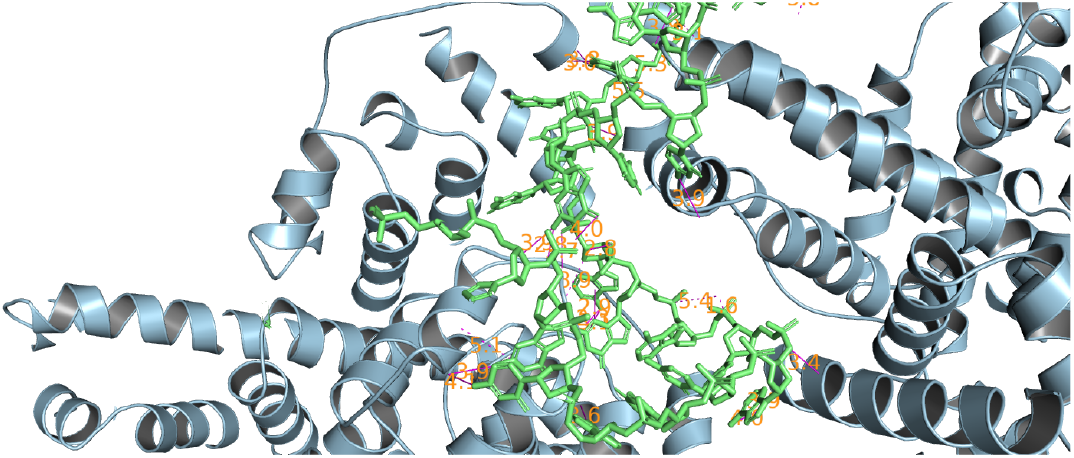
Molecular interaction between Ni_apt and BSA chains. Most relevant interactions take place in the domain III of both BSA chains. Figure made using Pymol.

The aptamers interaction sites on BSA were mapped as reported by Table 1. The top residues for docking were Lys, Glu and Lys Arg for R_apt and Ni_apt respectively. Specific amino acids are more likely to interact with specific nucleotide bases (Hossain, Kazi Amirul, et al 2023). Lysine especially represents the higher number of interactions in both dockings in accordance with Chan, Chen-Hui, et al. 2019; interestingly having a preference for adenine bases, contrasting the funding of Lumscombe et al (2001). The overall interaction network of Ni_apt aptamer suggests potentially stronger binding due to more hydrophobic interactions and hydrogen bonds, even if R_apt presents a possibly relevant pi-Cation interaction. Finally, threonine can form methyl-methyl contacts with thymine bases (Luscombe et al., 2001), represented only in the Ni_apt. Interestingly, there is a predicted binding to a guanidine base as well.

### 3.3 Measuring Protein Binding with BSA Aptamers and Fluorescent Intercalation Probes

#### 3.3.1 In vitro Binding of Aptamers with BSA

SYBR Green I is a very well characterised FIT probe for use against small molecules and proteins (Kong et al.2013). SYBR Green I and its binding to DNA ratio has been extensively characterised by Zipper et al. (2004) and has been optimised for characterising their binding behaviour against proteins to be used in a microarray (Cho et al 2006). This allows for very quick aptamer protein binding determination. For the aptamers used in this study SYBR Green I increases in fluorescence when the aptamer is bound to BSA (Figure 5) which corresponds to an increase in stacks where the fluorophore can intercalate. The BSA was titrated against the R_apt to obtain a Kd of 0.02 μM and a Bmax of 3.99 μM (Fig. 5, *purple curve*). The Ni_apt was found to have a Kd of 0.12 µM and a Bmax of 10.35 µM (Fig. 5, *aqua curve*). Both aptamers showed a large fluorescence change according to the BSA concentration in the solution.

**Figure 5.**
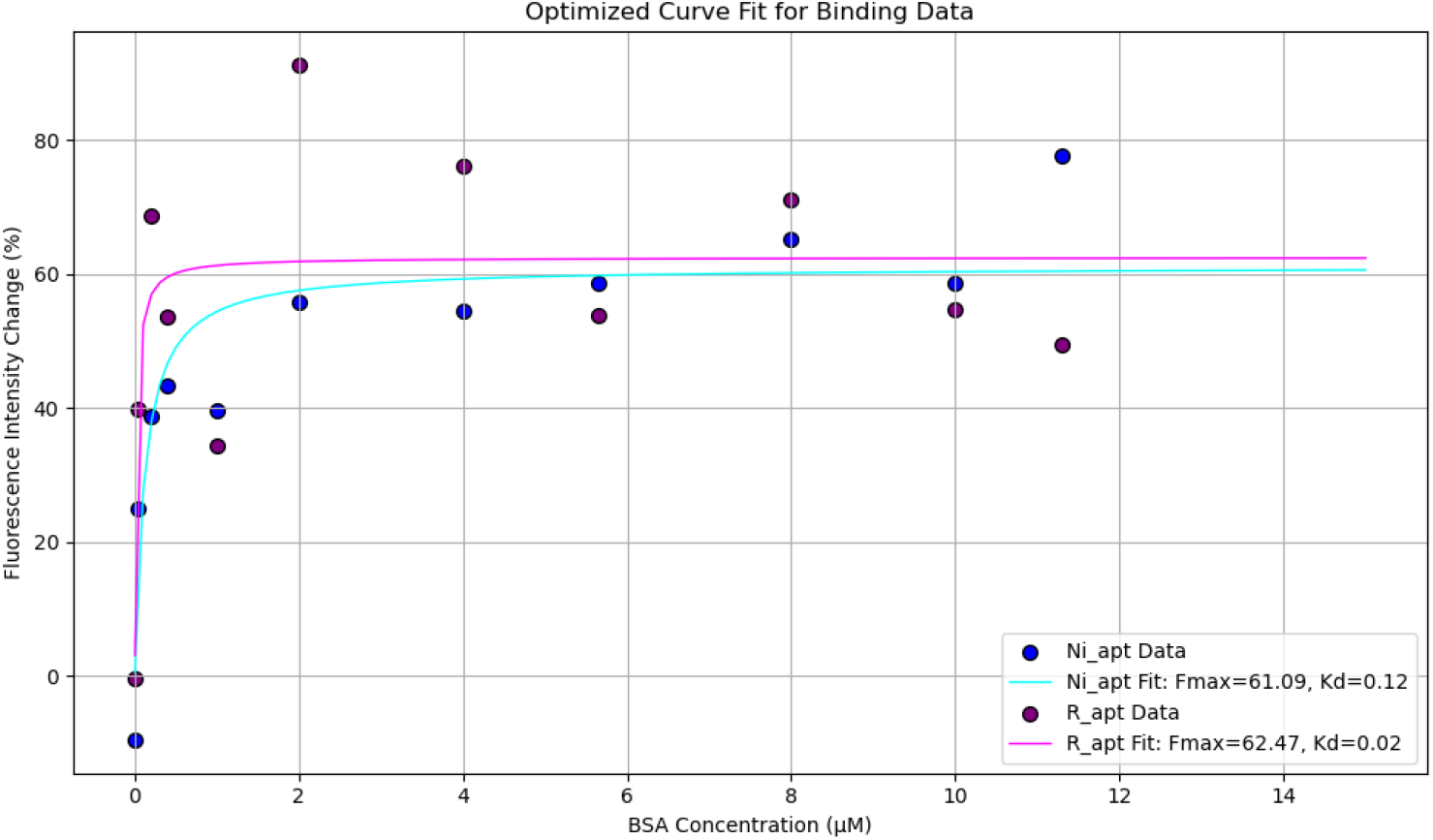
SYBR Green I Fluorescence Response to Ni_apt BSA Binding (aqua) Kd is 0.12 µM and R_apt (purple) Kd is 0.02 µM.

The R-squared of the two logarithmic fittings curves are respectively 0.8589 and 0.6094 for Ni_apt and R_apt. The binding interaction was confirmed by a correlation between rising BSA concentrations and increasing fluorescence intensity. Due to protocol changes SYBR Green I fluorescence overall decreased. No significant changes in fluorescence were observed with aptamers alone or in the buffer medium. Data points above the logarithmic curve fitting are due to possible introduction of small volume shifts. Larger intensity was detected when performing the measurements the same day of buffer preparation, with newly fresh solutions and performing a short incubation time.

### 3.4 Development of Aptamer-Based Affinity Chromatography

We designed the Ni_apt to have a polyA tail to give the same effects as a His-Tag linker. Ni-NTA magnetic beads are beneficial for pull down assays for purifying proteins of interest (Nastasijevic, 2008). As we found the aptamer, Ni_apt with the polyA tail to have a similar affinity for BSA, the Ni_spt would allow for selective binding from a dilute BSA solution, allowing up-concentration of this protein. Since the method used here is simple and results in similar binding both in silico and in vitro it can be used for a number of other protein biomarkers in the future (Perret, G., and E. Boschetti 2020).

To test the binding on the Ni-NTA magnetic beads we determined the starting concentration BSA in solution and determined the final quantity left in the wash solution using a Nanodrop absorbance spectrometer. After the bound material was washed it was determined that there was 79.3 % of the material left in the wash thus 20.7 % of the total BSA from the starting solution was bound to the beads (Fig. 6). However, these results could be improved in the future. Kökpinar et al (2011) found a 75% recovery when His-Tag proteins with anti-His-Tag aptamers. Likewise, human C-reactive protein was recovered with up to 87% recovery with an anti-CRP aptamer (Bernard et al 2015) and 45% recovery of activated protein C (Murphy et al 2003). The lower recovery we had was likely due to steric effects due to the large size of the BSA. We expect a higher recovery amount when a substantially larger number of beads complexed with aptamers are used. We are currently optimizing the conditions to obtain a higher recovery.

**Figure 6.**
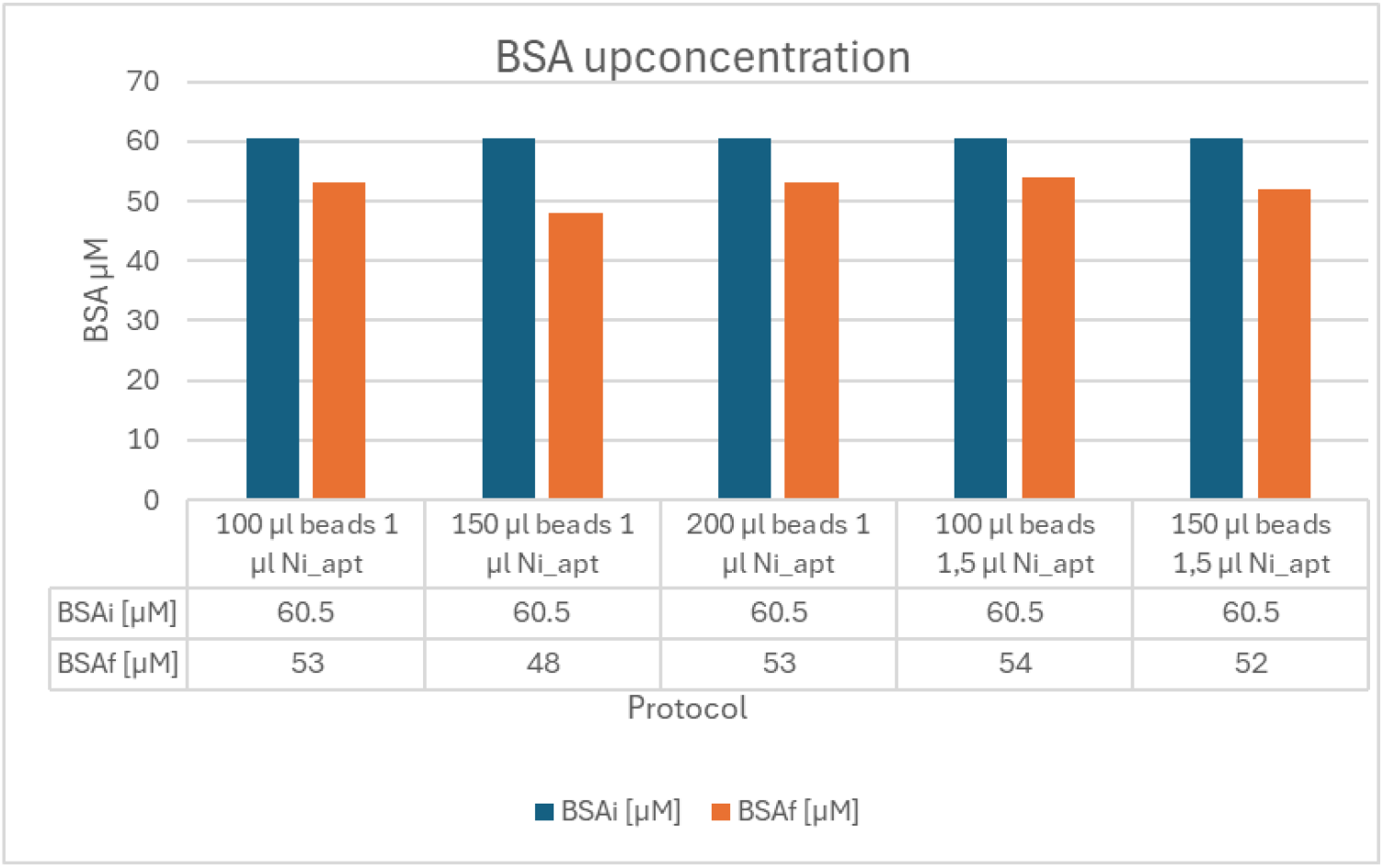
BSA up-concentration protocol results. Each bar of the graph represents different tested protocols with beads slurry and aptamer volume variations. The orange bars represent the elute concentration, the resulting variation from the starting BSA concentration (blue) represents the BSA bound to Ni_apt. BSAi is the initial concentration of BSA that was added to the aptamer with beads and BSAf is the final concentration found in the wash.

## 4. Discussion

A novel aptamer-based affinity chromatography method was explored to purify BSA using a BSA-specific binding DNA aptamer in this study. The focus was on validating the feasibility of the newly designed purification method and the binding efficiency of the specific aptamer using affinity chromatography. Through the SYBR Green I binding affinity experiments and tryptophan fluorescence experiments, we validated our hypothesis that aptamer-based affinity chromatography can serve as an efficient and cost-effective method for protein purification. Its binding efficiency and ability to maintain protein integrity and functionality throughout the process indicate that this method has the potential to be a viable alternative to traditional protein purification techniques.

### 4.1 Aptamer-BSA Binding Analysis

Using SYBR Green I fluorescence assays, we quantified the binding interactions between BSA and two aptamers: the standard aptamer and the polyA-tailed aptamer. McKeague et al (2015) found that SYBR Green I can compare to other techniques for finding the Kd. Additionally, SYBR Green I offers a very quick and easy approach to determining binding of aptamer–protein. The R_apt had a dissociation constant (Kd) of 0.02 µM, which is lower compared to the published results (Kd = 69.44 ± 7.60 nM) reported by Wang et al. (2019); indeed, different methods of theoretical Kd calculations were applied here from the published results. The Ni_apt had a Kd of 0.12 µM, slightly higher than the R_apt. Thus polyA tail modification did not seem to affect the aptamer binding to BSA and can be utilised for the Ni-NTA beads. Future experiments can focus on improving the binding affinity of the polyA-tailed aptamer with BSA to achieve a higher binding affinity for better BSA purification.

### 4.2 PyMOL/PLIP Analysis of Binding Sites

The PLIP, ligand interaction profiler, provided crucial information about the aptamer’s binding sites on BSA, further visualised with Pymol. We identified several interaction sites, particularly in domain I and III of BSA of both monomers, including: R143, Q579, T83 and R81 of monomer A and less closer interactions with the monomer B (cut-off 3 Å). This structural analysis is essential for understanding binding mechanics and guiding the development of future aptamers with greater specificity and affinity. Kuntip et al. (2021) found another aptamer that docked with BSA, Human Serum Albumin (HSA), and Canine Serum Albumin (CSA), and showed stronger and more specific binding to BSA. Both studies indicate that domain III of BSA has unique structural and physicochemical properties that enable high-affinity interactions. These findings suggest that future aptamer designs should target specific amino acids in domain III to improve binding affinity and specificity. By decreasing off-target effects and enhancing the effectiveness of protein purification procedures, future research based on this study may open the door for the development of more effective and targeted aptamers, which is essential for therapeutic applications. In conclusion, this research makes a significant contribution to the protein purification field by laying the groundwork for future advances in aptamer-based technology and its applications in biotechnology and medicine.

### 4.3 Ni-NTA Beads and Affinity Chromatography

Using Ni-NTA beads functionalized with the polyA-tailed aptamer was a key component in our method. We used the wash from the solution to determine BSA concentration with the Nanodrop. Our beads recovered 20.7% of BSA from a diluted solution at an initial concentration of 121 μM. This confirmed that the polyA tail did not interfere with the aptamer-BSA binding interactions while facilitating attachment to Ni-NTA beads.

The relationship between the number of magnetic beads and the concentration of Ni_apt bound to them depends on the binding capacity of the beads and the amount of aptamer available. To extract up to 20.7% of BSA from a concentrated solution we used 1 ul of Ni_apt (10 µm), gaining a dilution of 0.099 µM during the first incubation with the beads.

We then calculated the theoretical maximum amount of aptamer that can bind. This calculation considered the binding affinity of the beads for His-tagged proteins, as specified in the Cytiva documentation. The total binding capacity was determined by multiplying the bead volume by their binding capacity. Given a binding capacity of 50 mg/mL and a total bead volume of 0.025 mL, the total binding capacity was calculated as follows:

Total binding capacity=50 mg/mL×0.025 mL=1.25 mg

Thus, the theoretical maximum binding capacity was determined to be 1.25 mg.

Theoretical calculations show higher binding capacity of the beads leaving more free surface for Ni_apt to bind, however, the Nanodrop analysis revealed DNA impurities in the elute, opening the possibility of steric hindrance among aptamers even though the short aptamer sequence should cause minor steric hindrance than an His Tag tagged protein. In addition the binded BSA was expected to have more consistency. BSA is a high molecular weight protein that could interfere by steric hindrance and disrupt binding interactions between the aptamer.

Further studies can be conducted to relate the aptamer steric hindrance and the available binding surface of the beads to characterise the optimal ratio aptamer to magnetic beads. In future experiments, non-BSA-specific aptamers can be used as a control group helping to validate the specificity and efficacy of our affinity chromatography method. Protocol improvements for high diluted solutions of expensive compounds could possibly obtain high protein retention, avoiding steric hindrance.

## 5. Conclusion

In this study the Ni-NTA BSA aptamer was designed with a poly A tail that allows it to bind to Ni-NTA beads. Using a polyA tail allows a simple way to modify the aptamer without using a His-Tag and thus a streamlined aptamer can be developed purely with nucleic acids. We found that the polyA tail did not change its BSA interaction. Interestingly, when the same SYBR Green I assay was performed with this aptamer the Kd achieved was comparable. Thus the poly A tail did not affect the binding and could be used for our extraction method. Although there are some variations of the SYBR Green I assay used in this study, the assay provides a quick, easy and scalable way to perform Kd experiments. More studies will be conducted to optimise and perform reproducible results.

The aptamer used to bind to Ni-NTA beads was used in excess. By determining the surface binding of the aptamer to the beads we could increase the amount of protein bound to make an efficient purification system where 100% of the protein is captured on the beads. In this study, we did not elute the protein from the aptamer but several techniques such as enzymatic digestion could be used. BSA was used as an example protein to purify using aptamer Ni-NTA system. Since aptamers are versatile for binding to small molecules and proteins we can utilize this system for purifying a number of different small molecules and proteins. Recently, Dianox developed a large language model, AptaBERT, to predict and generate aptamers that can bind with high specificity and affinity. These aptamers generated with this model have been tested in vitro and shown to work in our lab with many disease relevant and environmentally relevant proteins. Utilizing AptaBERT, aptamers can be generated that can potentially be used to purify or upconcentrate a number of different proteins for research, diagnostic or therapeutic use.

